# Estimation of Interaction and Growth Parameters to Develop a Computational Model for Gut Bacteria

**DOI:** 10.1101/2023.03.22.533813

**Authors:** Himanshu Joshi, Meher K. Prakash

## Abstract

The relevance of gut bacterial balance to human health can not be overemphasized. The gut bacterial balance delicately relies on several factors inherent to the person as well as to the environment. As the volume of evidences for the gut bacterial influence on health and the clinical data on the variance of the bacterial population across cohorts continue to grow exponentially, it is important to develop a theoretical model for gut bacteria. In this work, we suggest a new computational method for estimating the interaction parameters from the cross-sectional data of bacterial abundances in a cohort, without requiring a longitudinal followup. We introduce a nutrient type based bacterial growth model and use the Monte Carlo approach to estimate the matrix of interaction parameters for the 14 major bacterial species. These parameters were used in a comprehensive first-level computational model we developed for the large intestine to understand the patterns of re-establishing balance with different nutrient types.

## 1 Introduction

The human gut is a complex community of about 2000 types of bacteria, adding up to over a trillion (10^12^) bacteria all together [1, 2]. The digestion of the food happens significantly in the gut, aided by the bacteria [3]. However, the role of gut bacteria is much more significant. Gut bacteria have a symbiotic relationship with the host [4]. The microbiota offers many benefits to the host through a range of physiological functions such as strengthening gut integrity or shaping the intestinal epithelium, harvesting energy from nutrients that are in the diet, protecting against pathogens, and regulating host immunity [5]. Gut bacteria confers resistance to a wide range of intestinal pathogens such as vancomycin-resistant *Enterococcous*, and *C*.*difficile* [6, 7].

The Human Genome Project was aimed at gaining a complete understanding of human genes, their function, and relevance for the diseases [8, 9, 10]. However, the genetic understanding of several diseases remained open as many genes were implicated in any disease. Following the genome project,”Human Microbiome Project” [11] initiated about 10 years ago to understand the symbiosis between human and their gut bacteria and to understand how microbes that reside in and on the human body, which are very much responsible for health and disease. Of these, the most significant are the gut bacteria, and a lot of data on the complex interactions of bacteria or their influences on the host are becoming available [12, 11, 4].

What is very interesting from many of the recent findings is that gut bacteria have a role that extends beyond the gut. Several recent metagenomics, next-generation DNA sequencing studies have established the connection between the intestinal microbial species compositions and extra-intestinal diseases [11, 4, 13]. Changes in bacterial compositions are being implicated in gastrointestinal diseases such as Crohn’s or metabolic disorders like obesity [14, 15, 16] and type-2 diabetes or extra-intestinal diseases like arthritis and even neurodegenerative disease like Parkinson’s [17].

The stability of the gut bacterial ecosystem is thus very important for the overall health of the host [12]. Changes in diet [18], excessive use of drugs for health care or in the food industry [19], or excessive growth of opportunistic bacteria [20] can change the compositions of these bacteria, which may have a serious effect on human health. Drug induced changes in the bacterial community can increase the risk of diseases like acute intestinal infections. Bacterial composition changes made by antibiotic drugs do not necessarily restore the initial composition of bacteria in the gut, and disturbed microbiota are often susceptible to pathogen invasions [21].

The physiological relevance of the balance gut bacteria draws attention to the importance of understanding their interactions among themselves with nutrients and drugs. Theoretical efforts are required to provide such insights and understanding. However, modeling the gut microbiota is complex. The literature on theoretical modeling of the culturable, pathogenic bacteria is fast expanding [22, 23, 24]. While a similar level of activity for developing gut bacterial models would be much welcome, the complexities of working with gut bacteria have limited the studies so far. The gut bacterial data is mainly from clinical studies rather than from culture studies; the inferential studies are to be based on the available data or with clinical trials. There have been several attempts at estimating the parameters that define bacterial interactions. However, while the method is simple, it has limitations in that the co-occurrence need not be the bacterial interaction, and the interaction is likely to be asymmetric, unlike the co-occurrence matrix [25, 26]. Other models used clinical data with longitudinal followup after a perturbation with a dietary change [27, 28, 6]. There are many other studies that use time series data to estimate interaction parameters [29, 30]. However, it is not easy to perform such longitudinal studies, and alternative approaches are required.

Added to the complexity of the interaction parameter estimation is that a clinically relevant model with interactions with diet or drugs also needs to be developed. To the best of our knowledge, there has been no comprehensive work that attempts to estimate the gut bacterial interactions and tries to correlate them to the clinical observations.

The aim of the work is to introduce a computational framework for comprehensive modeling of the gut bacterial populations and the nutrient. To achieve this, we combine various concepts, including bacterial growth models, nutrient association, bacterial interaction parameter estimation, and adaptation of the gut bacteria to dietary changes. The model should be a basis for gaining insights into the different scenarios of perturbations to the diet and bacteria.

## 2 Results and Discussion

### 2.1 Detailed Model

#### 2.1.1 Dynamics of the bacterial numbers and the nutrient

In general, a healthy adult gut is about 150 cm long, and it hosts around 1000 to 2000 strains of bacteria [31, 32]. However, as a practical first attempt, we modeled the 14 most abundant strains, listed in Table 3, from the phyla *Bacteroidetes* and *Firmicutes* found in a cohort of 97 adults [33, 16]. Interestingly, their growth with the different macronutrients in the diet is not the same [34]. In this study, we focus on the growth dynamics of these 14 types of bacteria with the four macronutrients, namely protein, fat, carbohydrates, and fiber.

The nutrients *C*_*j*_(*t*), *j* = 1 to 4 representing the concentrations of protein, fat, carbohydrate, and fiber, respectively, are assumed to continuously drift with a speed *v* along the large intestine, with the push from the muscles of the intestine. As the nutrients drift along the intestine, they are consumed by the bacteria for their own growth (*γ*_*b*_) as well as for that of the host (*γ*_*h*_).

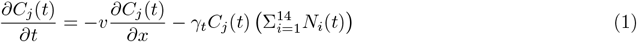

Simultaneously, we model the bacterial numbers *N*_*i*_(*x, t*) along large intestine as

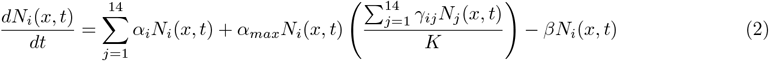

where, *N*_*i*_(*x, t*) are the population densities of the 14 bacteria at position *x* and time *t*, respectively. The three terms on the right-hand side of the equation (1), described in detail below, represent the contributions from the growth and multiplication, competition with the same as well as with other species, and death, respectively.

##### Growth

We model the growth rate of the bacteria *i* with the all the nutrients *j* as

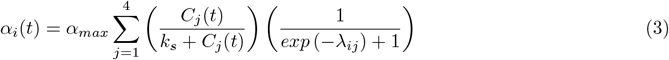

In equation (3), *α*_*max*_ which is the maximum growth rate of each bacteria with the most appropriate nutrient type. We assume that in the exponential phase doubling time of the bacteria is 25 minutes and use the corresponding *α* _*max*_= 0.0007 per second. The second factor 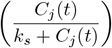, similar in spirit to the Michaelis-Menten kinetics, ensures the saturation of bacterial growth rate at high concentrations *C*_*j*_ of the nutrient *j. k*_*s*_ is the nutrient concentration when the growth rate is half its maximum value. In our model, we assume *k*_*s*_ = 5.0 *g/cm*; since the gut is modeled at high nutrient concentration, the results are not sensitive to *k*_*s*_. And the third factor modifies the rate using the bacteria-nutrient association factor *λ*_*ij*_. Since the bacterial growth is positive regardless of whether the association rate is negative or positive, we used this exponential factor. From the experimentally determined [34] association of dietary components protein, fat, carbohydrate, and fiber with bacterial strains for 14 most abundant bacterial strains, values of *λ*_*ij*_ are given in Table 3.

##### Competition

Despite the availability of the typically represents the capacity of the medium to support the bacterial population. In this, we consider the colon as a 1*D* linear structure, with an implicit radius of crosssection of 3 *cm*. Assuming 10^12^ bacteria over the 150 *cm* length of the colon, for the spatial capacity *K*, we use a one-dimensional linear density per each of the 1 *cm* segments, *K* = 10^8^. Constrained by this capacity of the colon, the different bacteria compete among themselves.

##### Death

In this model, we are using a death rate (*β*) equal to 9 × 10^*−*7^ per sec. To verify if this value of *β* is physically meaningful, we gave some nutrient concentration to bacteria and let them grow for 24 hours. When we plotted bacterial density versus time, we obtained the following growth curves for bacteria. By comparing these logistic growth curves with the experimental logistic growth curves, we believe that we are using a reasonable value for the death rate.

#### 2.1.2 Cases with and without spatial component

The detailed model discussed above includes the growth of bacteria at every stage of the drift of nutrients. However, for calibrating parameters, whether it is the growth rate or the interaction parameters, studying the growth dynamics at a single position or studying them with the detailed model did not make any difference. Hence, for parameter estimations, we use the model with no spatial variation of the nutrients, dropping the index *x*.

#### 2.1.3 Relevance of data used in the model

To model the gut bacterial population, we need to estimate the growth rate of gut bacteria with diet nutrient environment and interaction parameters of gut bacteria with each other. To estimate the growth and interaction parameters, we used the following studies. The observational clinical studies on 98 individuals which focus on the effect of diet on gut bacteria using inferential techniques [34]. They conclude that the gut bacteria get affected by the diet within a day and quantify the association of gut bacterial strains with the macronutrients present in the diet, which we used to model the growth rate of bacteria in different macronutrient environments. In 2014, another study on the effect of diet on gut bacteria was published [35]. This work aims to check how fast diet can alter gut bacteria and whether this is reversible. In this work, the author concludes that plant- or animal-based diets change gut bacteria significantly in a week, and this effect is reversible. But they do not provide any data on the association between macronutrients present in the diet and gut bacteria. Apart from these two papers, we could not find any relevant study that tried to quantify gut bacteria’s association with diet clinically or experimentally. In this work, we use longitudinal bacterial abundance data for inferential estimation of bacterial interaction parameters. To quantify these interactions, we used the base abundance of gut bacteria in healthy humans [33]. This data is relevant for this study because this gives the bacterial abundance of 109 bacterial species in 97 healthy human samples.

### 2.2 Numerical solution for the model

The above equations 1 and 2 involve deterministic dynamics of bacterial growth with nutrient, *with yet to be determined* parameters. Thus, solving the above set of equations is required simultaneously to use a deterministic approach for the population and a Monte Carlo method for the parameters. The parameters are initially guessed randomly and updated, while the bacterial numbers follow the deterministic solution. Some of the parameter values used in the above equation are all listed in Table 1. Other parameters (*α*_*ij*_) need to be estimated. Simulations over a 4 months span were performed in response to the dietary defined in Table 2. After choosing a dietary plan, we gave a new pulse of nutrients every 6 hour for 4 months. The patterns of how the nutrients are consumed over the 6 hours are shown in Figure 7. The results of giving different dietary plans (Table 2) with a nutritional pulse of 500 *kcal* every 6 hours (corresponding to 2000 *kcal*/day) were studied in this work.

**Table 1:** List of parameters. The table gives the different parameters used in Equations (1), (2), and (3) and the sources where applicable.

| Parameter | Symbol | Value | Source |
| --- | --- | --- | --- |
| Nutrient consumption for bacteria metabolism | $\gamma_b$ | 0.2fg per bacteria per sec | [39, 40] |
| Nutrient consumption for host | $\gamma_h$ | 0.2pg per bacteria per sec | Assumed |
| Maximum growth rate | $\alpha$ | 0.0007 per sec | [41] |
| Death rate | $\beta$ | $\sim 10^{-6}$ per sec | Estimated [42] |
| Mass of the bacteria | $m$ | $\sim 10^{-12}$ gram | [43] |
| Nutrient Drift | $v$ | 0.008 cm per sec | Assumed |
| Michaelis-Menton constant | $k_s$ | 5.0 gram | Assumed |

**Table 2:** Nutrient composition. We used these dieting plans for four months. 1 *g* of protein contains 4 *kcal* of energy, 1 *g* of carbohydrate contains 4 *kcal* of energy, 1 *g* of fat contains 9 *kcal* of energy and 1 *g* of fiber contains 2 *kcal* of energy. So, each diet pulse contains approximately 500.0 *kcal* of energy.

| Diet/Nutrient | Protein (g) | Carbohydrate (g) | Fat (g) | Fiber (g) |
| --- | --- | --- | --- | --- |
| Balanced diet | 12.5 | 12.5 | 5.6 | 25.0 |
| High protein | 35.0 | 5.0 | 2.2 | 10 |
| High carbohydrate | 5.0 | 35.0 | 2.2 | 10 |
| High fat | 5.0 | 5.0 | 15.6 | 10 |
| High carbohydrate | 5.0 | 5.0 | 2.2 | 70 |

**Table 3:**
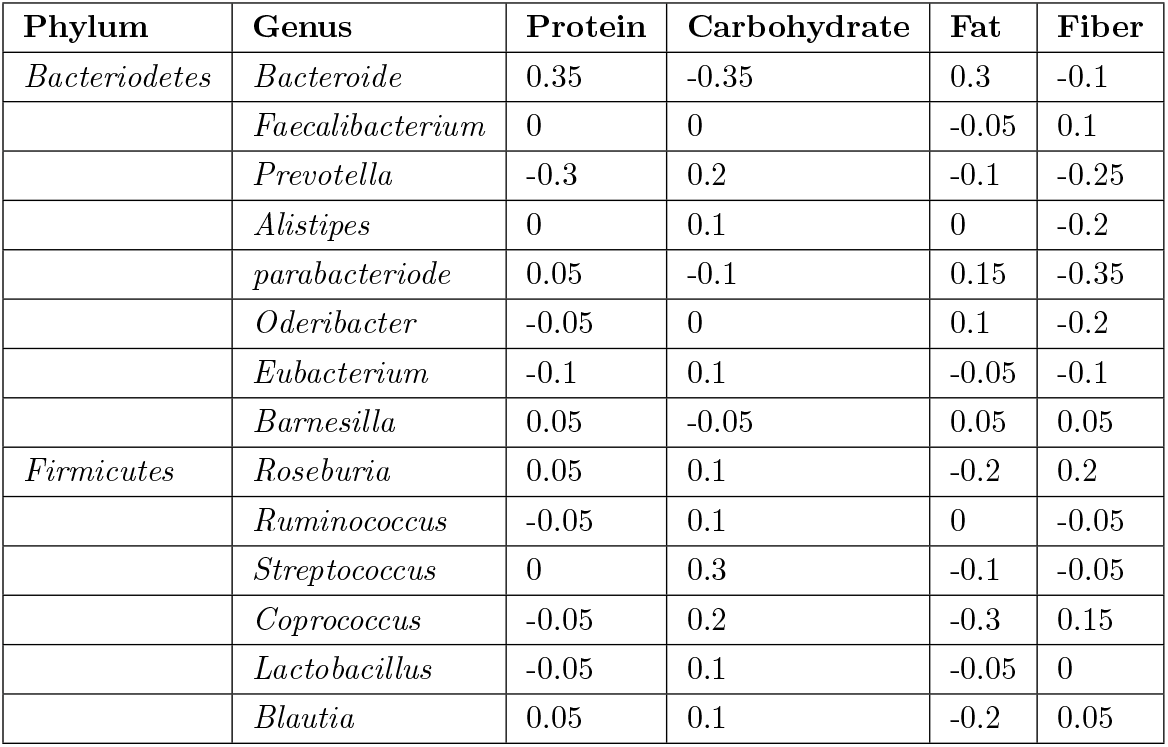
Bacteria-nutrient association. The association of the different strains of bacteria with the different macronutrients are obtained from the literature. These values are used as *λ*_*ij*_ in our model [34].

We numerically solved the above mentioned differential equation, which simultaneously handles the drift and absorption of the nutrients as well as the growth in the bacteria with our implementation in FORTRAN (code available, see **Additional Information**). Equations (1) and (2) were simultaneously solved using the forward-difference method. The updates during the simulation were guided by tracking the root mean square error (*RMSE*) of the bacterial populations as in Equation (4).

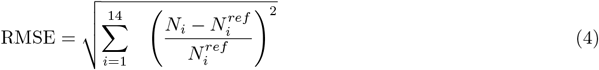

In equation (4), *N*_*i*_ and 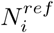 are the predicted and reference abundances for the *i*^*th*^ bacteria. To find the elements of the bacterial species co-occurrence matrix, as in Figure 2(f), Pearson correlation between bacterial species’ abundance from the 97 samples [33] was calculated. To quantify the association of bacteria with the nutrient type (Figure 10), we increase the one nutrient and see quantified the change in the bacterial population due to this diet plan.

### 2.3 Interaction parameters

#### 2.3.1 Algorithm for parameter estimation

We modeled the 14 most abundant bacteria in the human large intestine using the clinical data as a benchmark reference [33] and estimated the 196 asymmetric interactions among them. An initial population of 10^5^ of each bacterial strain and interaction parameters randomly chosen from the [-1, 1] range were used to initialize the population growth dynamics. A schematic of the parameter estimation algorithm is shown in Figure 1. The algorithm follows a deterministic Lotka-Voltera dynamics for modeling the bacterial growth with the 196 interaction parameters, which are the *current estimates* and need to be updated. The update of the estimates for the interaction parameter is guided by tracking the root mean square error (*RMSE*) of the bacterial populations as in Equation (4).

**Figure 1:**
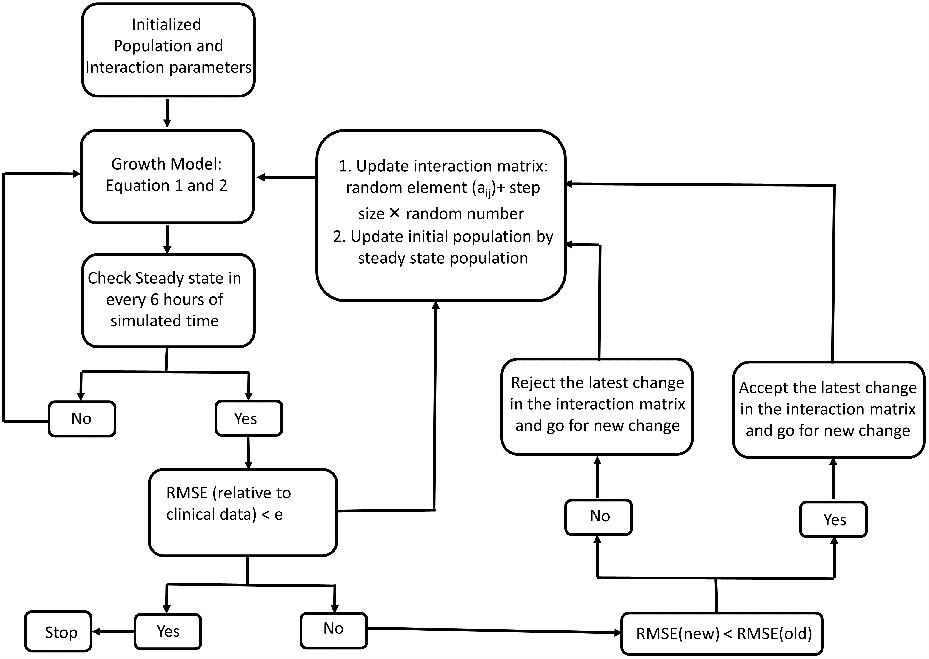
Parameter estimation procedure. The flowchart of the various steps involved in the interaction parameter estimation is shown. The procedure simultaneously uses deterministic equations for tracking the bacterial growth numbers, as well as updates the interaction parameters using a Monte Carlo approach. The population from clinical samples serves as a reference.

In the implementation of our algorithm, the same definition of *RMSE* from Equation (4) is used in two stages of calculation with two different reference populations. Using the abundances from the previous time-step in the dynamics as a reference, the approach to steady state condition for a *given* estimate of the parameters is monitored while using the clinical abundances as the reference, the convergence or the need for an update of the parameter estimates is monitored.

After the saturation, unless the populations converge to the clinical distribution, the next iteration step is performed by randomly choosing the interaction element to be updated by an increment which is also randomly selected. The deterministic dynamics are continued with this updated parameter, and if the *RMSE* relative to the clinical data improves, the update is accepted. Otherwise, the procedure is repeated with another randomly chosen interaction element. The iteration steps continue until the bacterial numbers correspond to the clinical data with no further improvement in *RMSE*. We also checked if, in a Metropolis sense, accepting the updates which worsen the *RMSE* with a small probability of 0.1 will change the results. Occasionally accepting the worse updates did not affect our results (data not shown), and it was not used in further analyses to keep the implementation simple. The various relevant metrics were tracked through the simulation with a graphical representation of the results, as shown in Figure 2.

**Figure 2:**
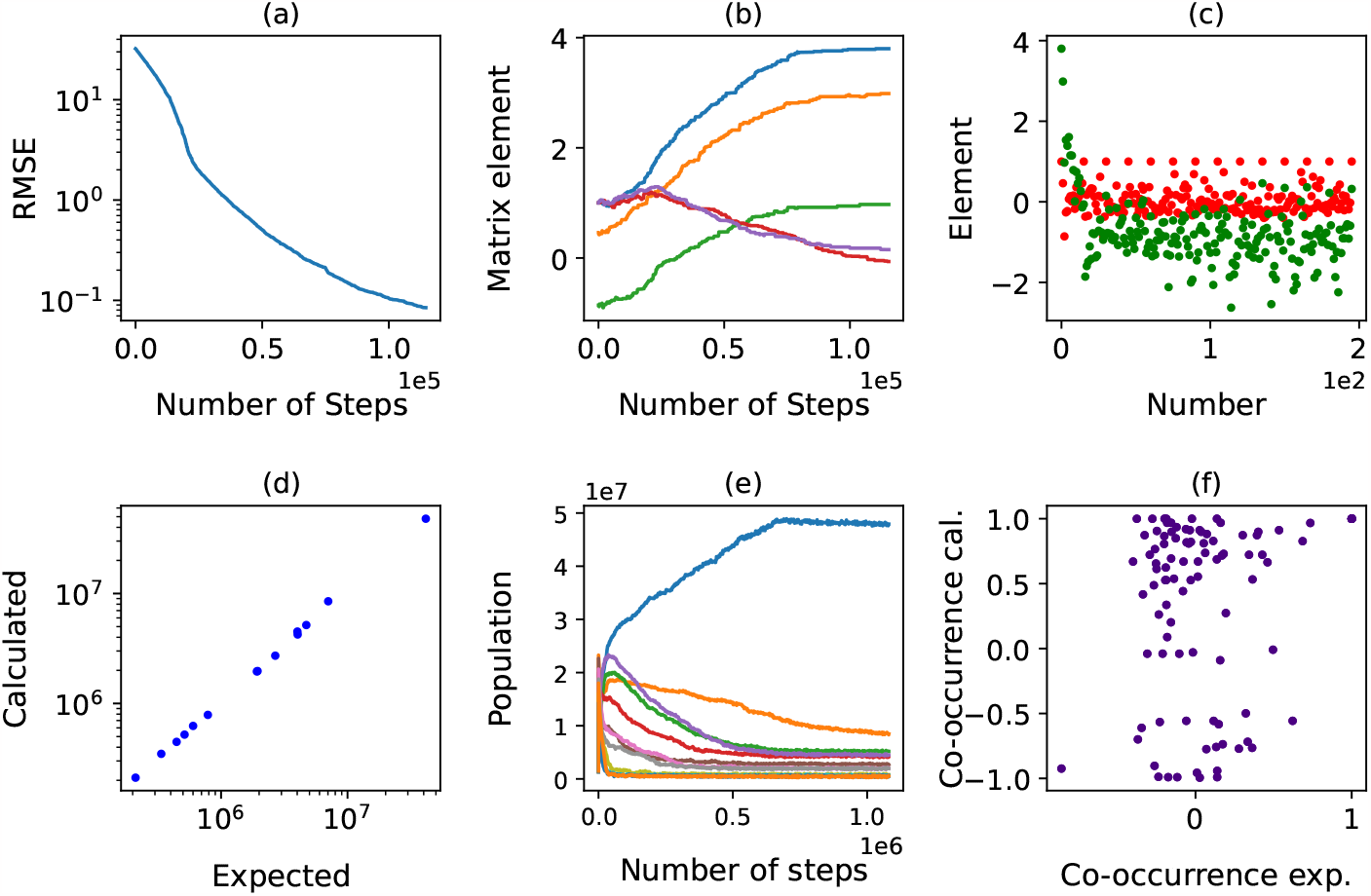
Parameter estimation dashboard. To track the quality of the estimated parameters, several metrics were tracked. (a) RMSE of the bacterial abundance with respect to expected abundance, (b) a few elements of the interaction matrix as they progress through the iterations, (c) the 196 interaction elements at the start of the iterations (red) and end of the iterations (green) between all pairs of the 14 enterotypes, (d) Expected and predicted abundance of bacteria, (e) the pattern of variation of the bacterial populations over the iterations, (f) comparison fo the co-occurrence from the 97 samples with the estimates made using a perturbative approach.

#### 2.3.2 Interaction matrix structure

The interaction parameter matrix for all the 196 pairs, including the self-interactions, is shown in Figure 3. In summary, the most dominant parameters are the negative self-interactions along the diagonal and the positive interaction of all bacteria with the most abundant bacteria. Further, as one may also guess from this interaction of the dominant strain, the interaction matrix is asymmetric. These asymmetries may be expected, as one bacterial strain may be effective in enhancing or diminishing the growth of another strain, while the reverse may not be true. A symmetric co-occurrence matrix as a guess for the interaction matrix will limit such possibilities, in addition to other limitations.

**Figure 3:**
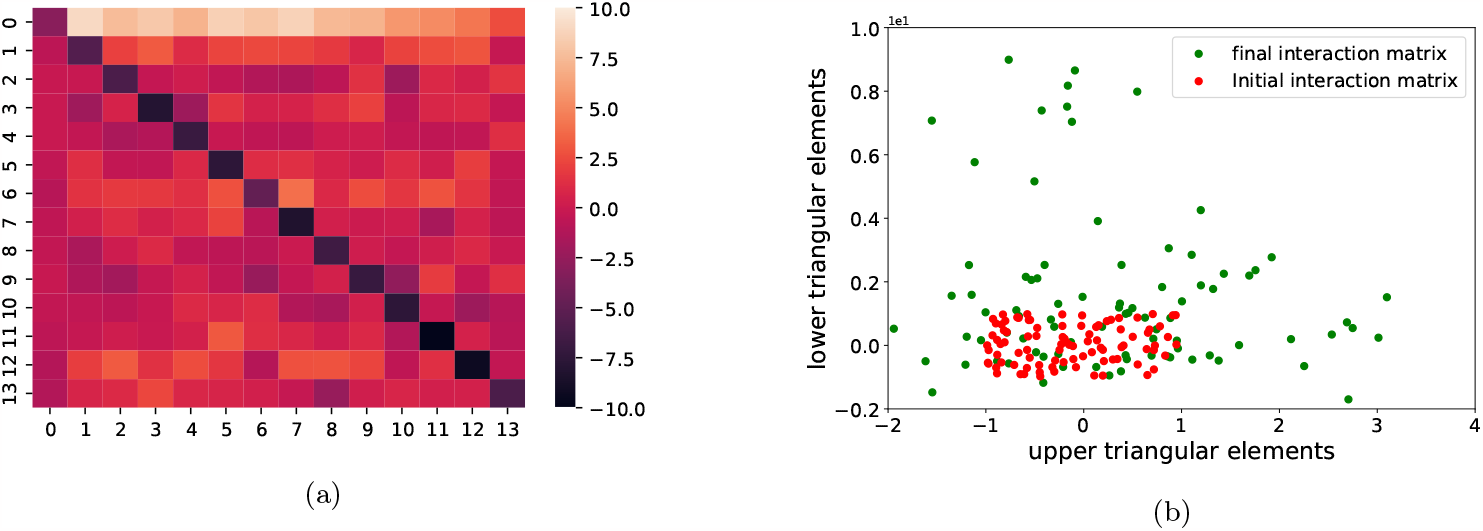
Pattern of interaction matrix elements. The estimated parameters are plotted to show (a) the range of variability and (b) the inherent asymmetry in interactions by plotting *α*_*ij*_ against *α*_*ji*_. The colorbar indicates the scale in (a).

#### 2.3.3 Relative insensitivity to the initial guess within a given range

The initial guess for the interaction parameters was generated by random number generation in the range [-1, 1] for all 196 parameters. The estimation procedure was repeated for several initial seeds that were used to generate the initial guesses for the interaction parameters. The estimated parameters were relatively insensitive to these initial conditions, a demonstration of which is shown in Figure 4. Also, different updating protocols, such as with different increment sizes, and also in the spirit of the Metropolis criterion, iterations that made the RMSE worse were also accepted with a small probability. The parameters were comparable.

**Figure 4:**
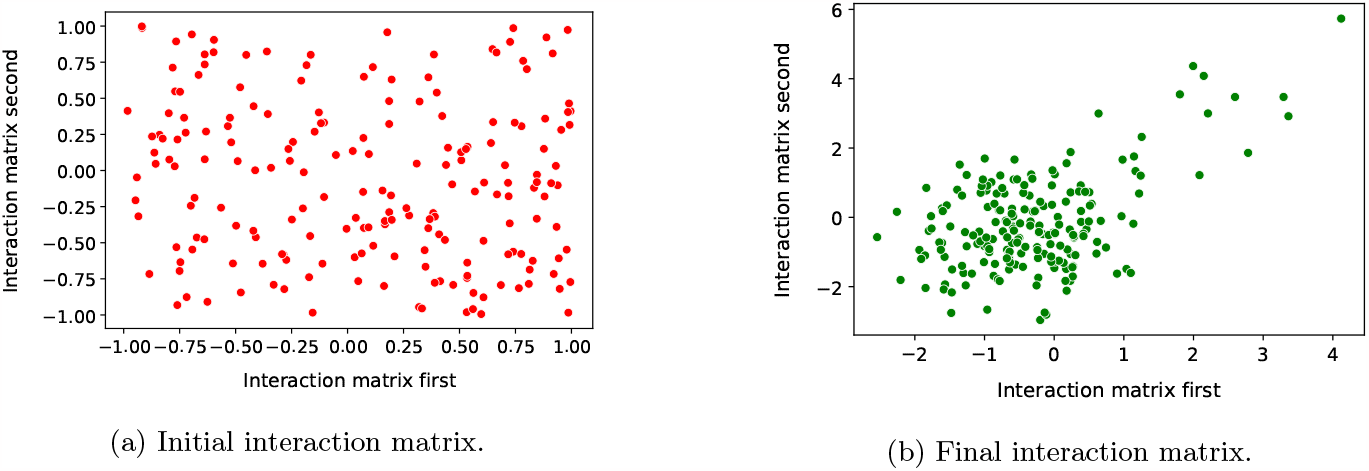
Invariance to random inital guess. The figure illustrates (a) two completely different random set of 196 initial interaction parameters, drawn randomly from [-1, 1] result in (b) similar final interaction parameters.

#### 2.3.4 Effect of the range of initial guess

However, the choice of the range [-1, 1] for the initial parameter guess was arbitrary. We repeated the estimation procedure by extending this range up to [-12, 12]. The bacterial populations converged to the clinical reference values in all these cases. While the convergence to the clinical values is, in theory, not guaranteed, it was a criterion for the estimation to be completed. The estimated parameters differ by a scaling factor, as shown in Figure 14. We distinguish the different estimates of the interaction parameters using an alternative metric not used in the estimation procedure. The rates at which the bacterial population reaches a new equilibrium following a perturbation were calculated. Figure 5 shows a specific example of one bacterial strain reaching equilibrium following a perturbation. The time to equilibration varies with the range of numbers used for the initial guess and saturates very quickly. Using additional information from the clinical data, a new equilibrium is established within a couple of weeks after dietary perturbations. We chose to work with the parameters obtained from the initial guess in the range [-1, 1].

**Figure 5:**
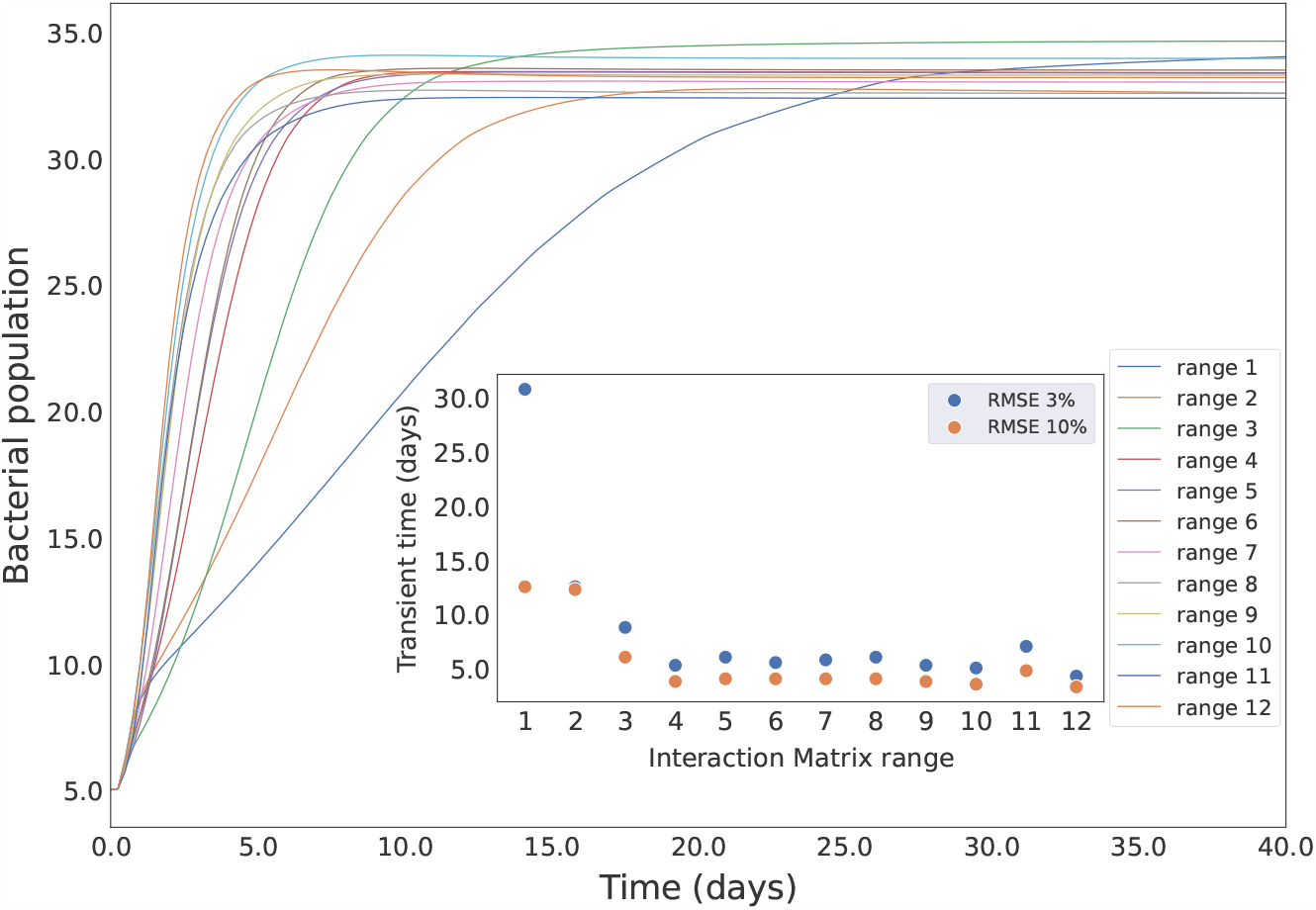
Time to equilibration. The initial guess for the interaction parameters is randomly chosen from a range of numbers. As the range increased from [-1, 1] to [-12, 12], factor 1 to 12, the rate at which the population equilibrated varied (inset). However, this did not make a difference at the clinical observation level.

#### 2.3.5 Variation across the multiple samples

The clinical data we used in the analysis was from 97 different patients [33]. We followed two alternative approaches to see the reliability of predictions. The first approach used the average numbers for each bacterial species to estimate the parameters. In a separate analysis, the estimation was repeated for all 97 samples. The average parameters obtained from this procedure were similar to those from the first approach Figure 6. The correlation between the two approaches shows the robustness of the estimates. While repeating the calculations with the individual samples was computationally expensive, it also suggests that such efforts may not be required, especially when dealing with very large cohort data.

**Figure 6:**
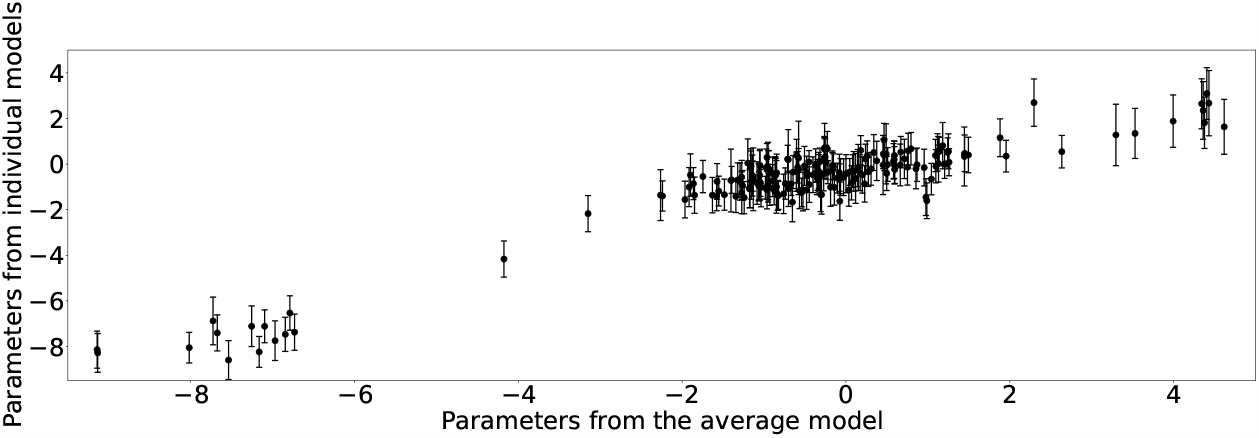
Ensemble vs individidual samples. The parameters estimated using the averaged bacterial populations as a reference were compared with those estimated from *each* individual patient sample. The mean and variance of the parameters are shown for the latter. The correlation between the two procedures is shown.

#### 2.3.6 Parameter estimation quality

Clearly, the estimation of parameters does not fall in the conventional notion where there are sufficient observations to estimate the parameters. Challenges such as this occurs commonly in many biological studies, for example, in studying microarrays [36], and they are typically dealt with in the framework of high-dimensionality statistics [37]. In the present work, we followed a Monte Carlo approach to establish the robustness of the estimates. It must also be noted that even the bacterial populations themselves are quite variable within a cohort of the adult population. Thus, the parameter estimation may be considered robust for modeling the colon. In addition, the present approach avoids the need for bacterial profiling from stool samples with follow-up on a cohort with longitudinal studies. We use these parameter estimates in the gut bacterial model.

### 2.4 Modelling the distribution of bacteria in large intestine

In modeling the gut bacterial distribution in the large intestine, there are a few comparisons that could be made with the clinical outcomes - the time to adapt to the dietary changes, which also serves as a validation and a basis for selecting the parameters; the nutrient-diet type association constants.

#### 2.4.1 Simultaneous changes in nutrient and bacteria

After the parameter estimations, the detailed model with simultaneous equations for the nutrient and the bacterial numbers is solved. The nutrient drifts as well as gets consumed with time (Figure 7). Figure 8 shows the distribution of bacteria in the large intestine. Towards the end of the intestine, the bacterial abundances are decreased because the nutrient availability is reduced.

**Figure 7:**
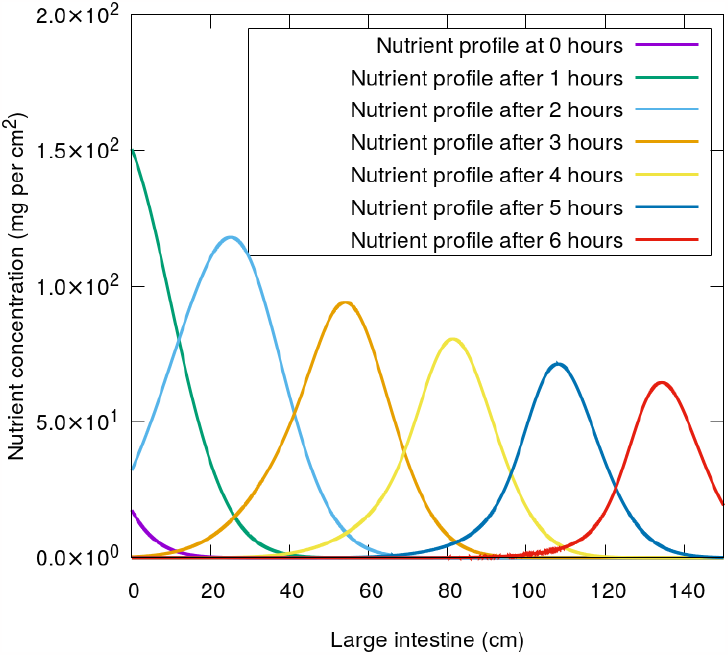
Nutrient profile along the large intestine. The large intestine is modeled as a 150 *cm* long tube, and the nutrients are assumed to be passing every 6 hour interval. The figure shows the nutrient profile along the 150 *cm* colon in a single 6 hour pulse.

**Figure 8:**
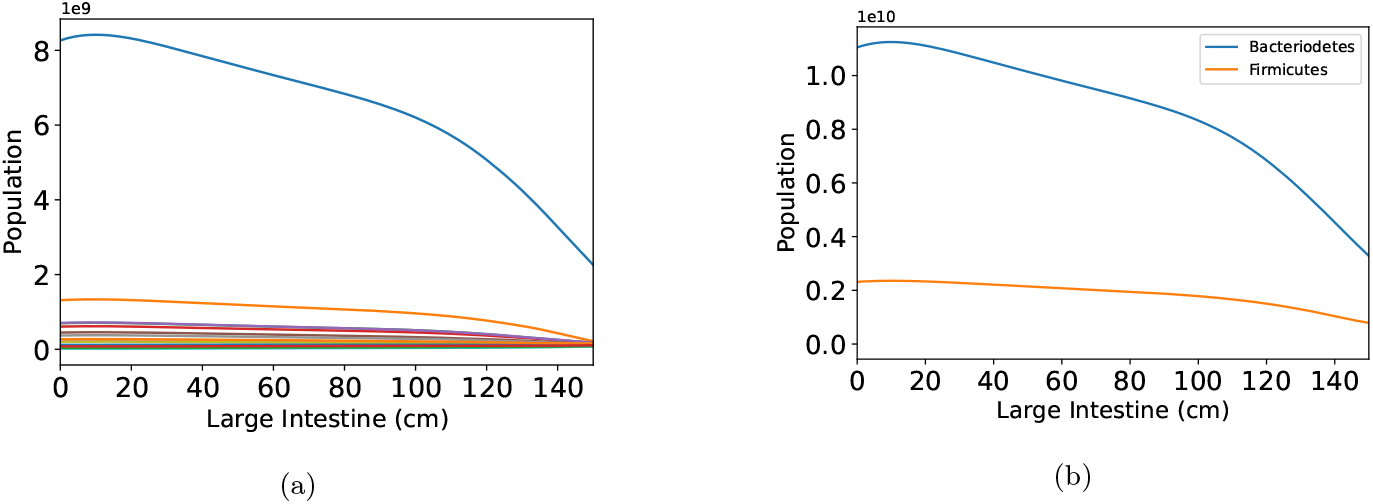
Spatial profile of bacteria. Distribution of the bacterial population in the large intestine for (a) each of the individual 14 bacterial types and (b) a regrouping of them as *Bacterioride* and *Firmicutes* phyla.

#### 2.4.2 Estimating the bacterial association with diet

As shown in Figure 9 the relative abundance of bacteria in the different diets has a significance effect on gut bacteria which are low in number. Bacterial abundance from the experimental studies matches with the high fiber diet and high carbohydrate diet.

**Figure 9:**
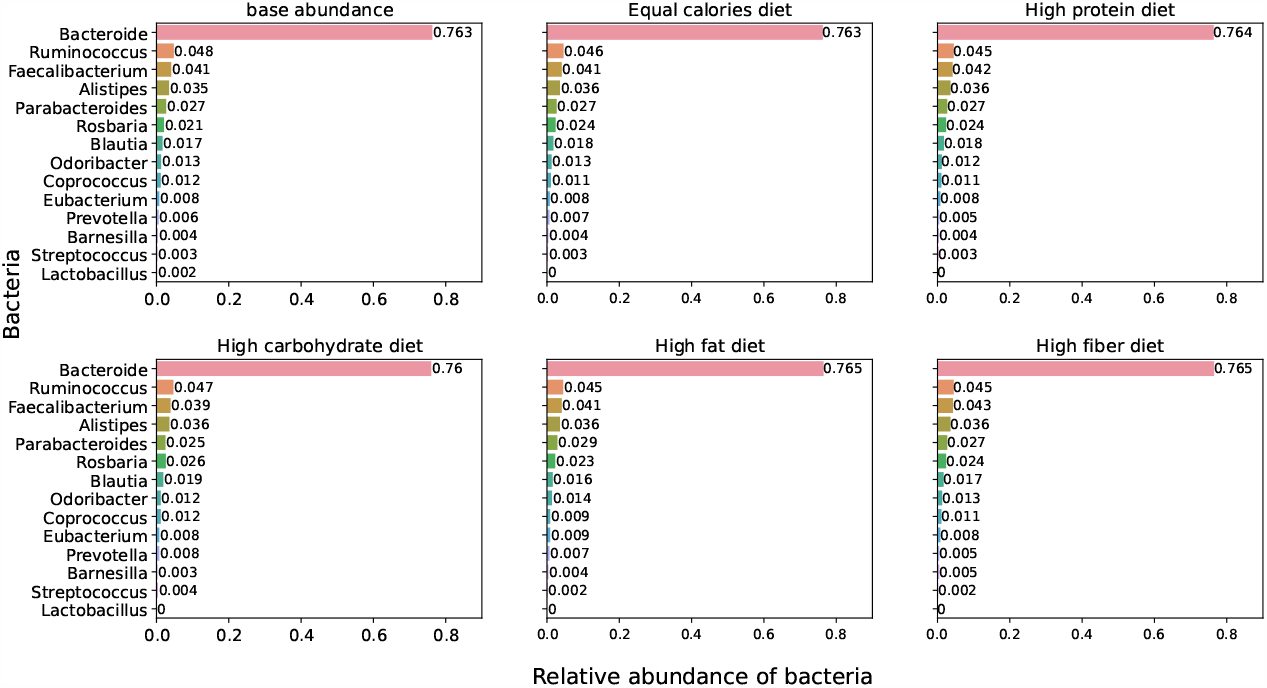
Relative abundance with different diets. The figure shows the calculated abundance of different bacteria under each diet plan. Notice the relative abundance of *Bacteroides*, which is almost constant except in the case of a high carbohydrate diet. A closer view of all the non-abundant bacteria is in Figure 15.

### 2.5 Comparisons with clinical data

#### 2.5.1 Association of bacteria with diet

The changes in the bacterial population with different diet types were monitored and compared to the clinical samples. This was done by monitoring the differences in each strain as the diet was changed from the equal calorie diet to a specific diet, such as the high protein diet. The changes are then checked relative to the bacterial association with diet from the literature. Figure 10 suggests that the association with diet is captured adequately. While this may seem obvious as the association enters our model as *λ*_*ij*_, the exponential factor we assumed with *λ*_*ij*_, the different stages of calculation after using the growth rates, and the difference in how it is estimated in clinical literature all suggest that this correlation is indeed non-trivial.

**Figure 10:**
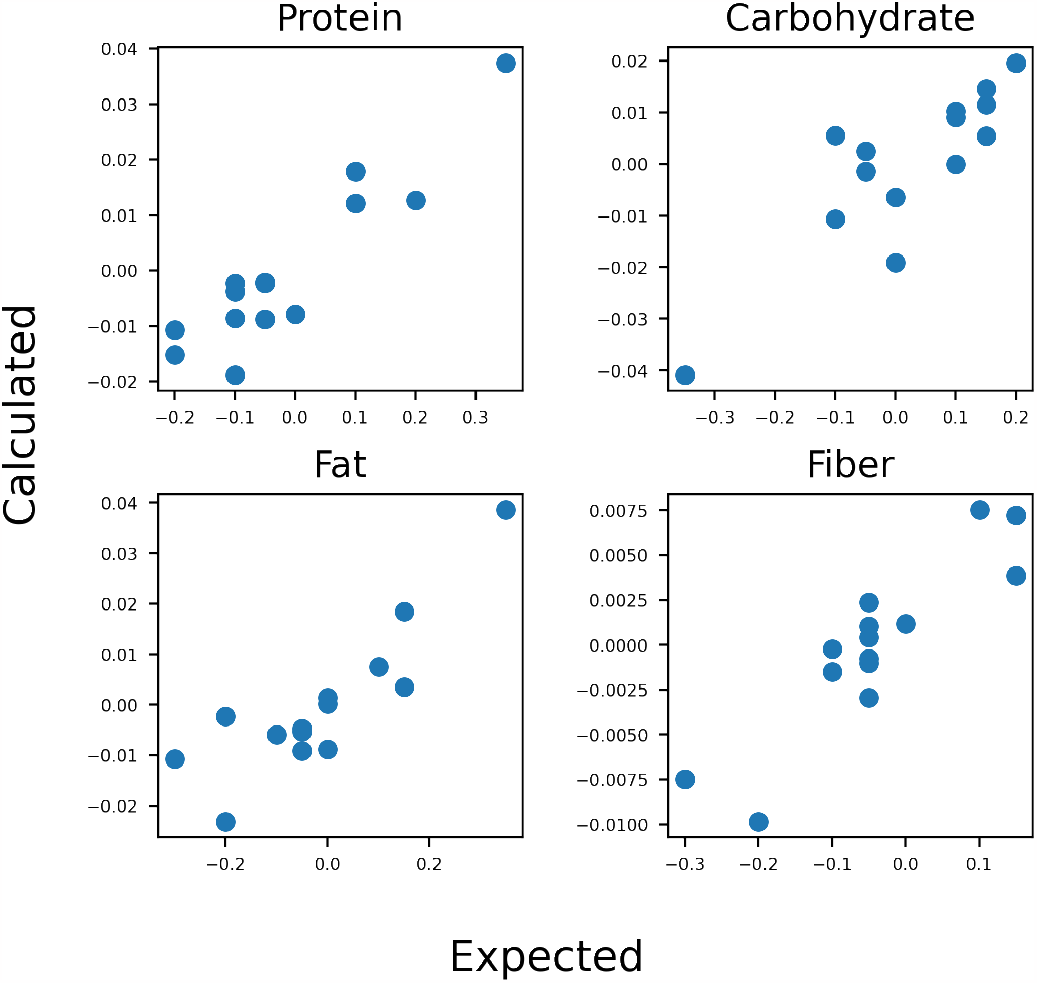
Diet-Bacteria association. The association of bacteria with the different diets is estimated by measuring the difference in the bacterial numbers obtained with each special diet relative to the balanced diet. These are compared with the association constants *λ*_*ij*_ from the literature.

#### 2.5.2 Re-equilibration time

Following any perturbation in the diet, the bacterial populations find a new equilibrium over time. The time to equilibration was comparable whether we studied the growth of bacterial populations starting from small seed populations or examined the time for re-equilibration. Figure 11 shows the equilibration patterns in the former case, starting from a seed population. The time to equilibration in these analyses was around 12 days. While the order of magnitude of the equilibration time was set by Figure 5.

**Figure 11:**
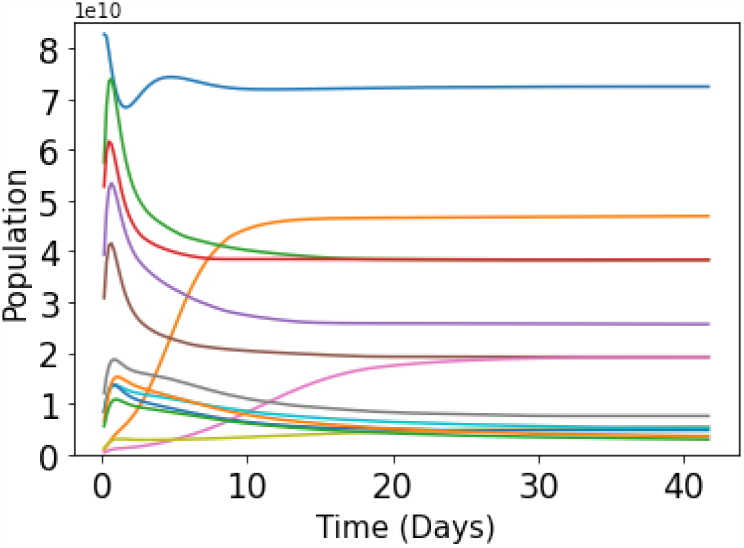
Time to equilibration. The above graphic shows how the population of the different bacterial strains re-equilibrates when an equal calories diet is given as a perturbation. The equilibration dynamics were fit to an exponential model to obtain the equilibration times.

**Figure 12:**
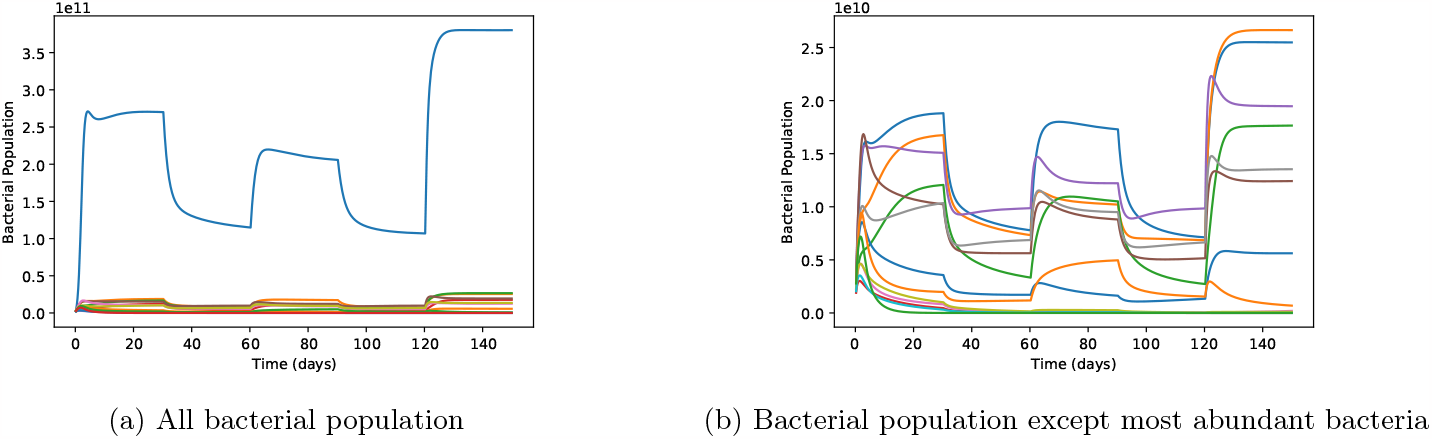
Simulated changes in bacteria with dietary plans. The effect of different dietary plans: equal calories (1−30 days), high carbohydrate (31−60 days), high protein (61−90 days), high fat (91−120 days), and high fiber (121 − 150 days) diets were calculated. The representation shows (a) all bacterial species considered in the model and (b) all other than the most abundant bacteria.

**Figure 13:**
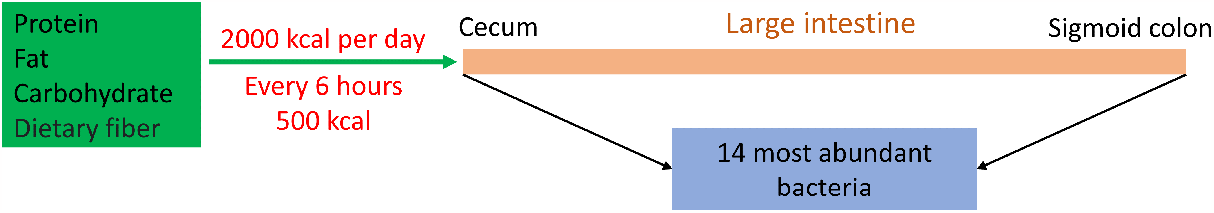
Gut model schematic. The schematic shows how the assumptions in the model. The 2000 *kcal* adult nutrition is assumed to be ingested in 4 pulses at 6 hour intervals. Different combinations of the macronutrients enter the large intestine after a pre-digestion of about 60% in the stomach and small intestine. The large intestine is assumed to be a tube that is 150 cm long and has a 4.5 cm diameter and receives 200 *kcal* from each pulse of 500 *kcal*. Our numerical solution uses a discretization of the 150 *cm* into 1 *cm* long segments.

**Figure 14:**
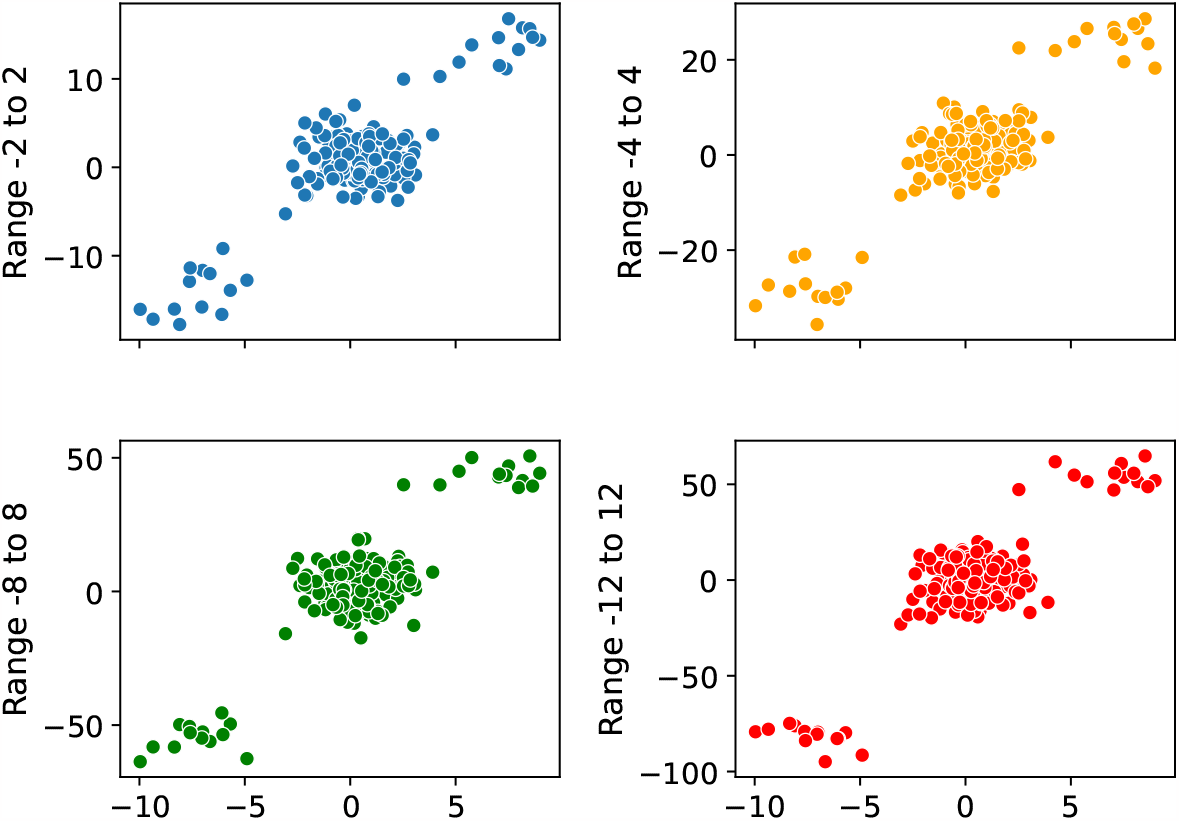
Effect of the range of initial guess. The plot shows the 196 parameters estimated using initial random guesses from the range [-1, 1] (x-axis) and their relation to the same parameters when estimated using initial random guesses from the ranges [-2, 2], [-4, 4], [-8, 8] and [-12, 12] respectively. The self-interaction terms and the interactions with the most abundant bacteria scale with the choice of the range. We use clinical data on re-equilibration times to choose the range [-1,1] as the most appropriate one.

**Figure 15:**
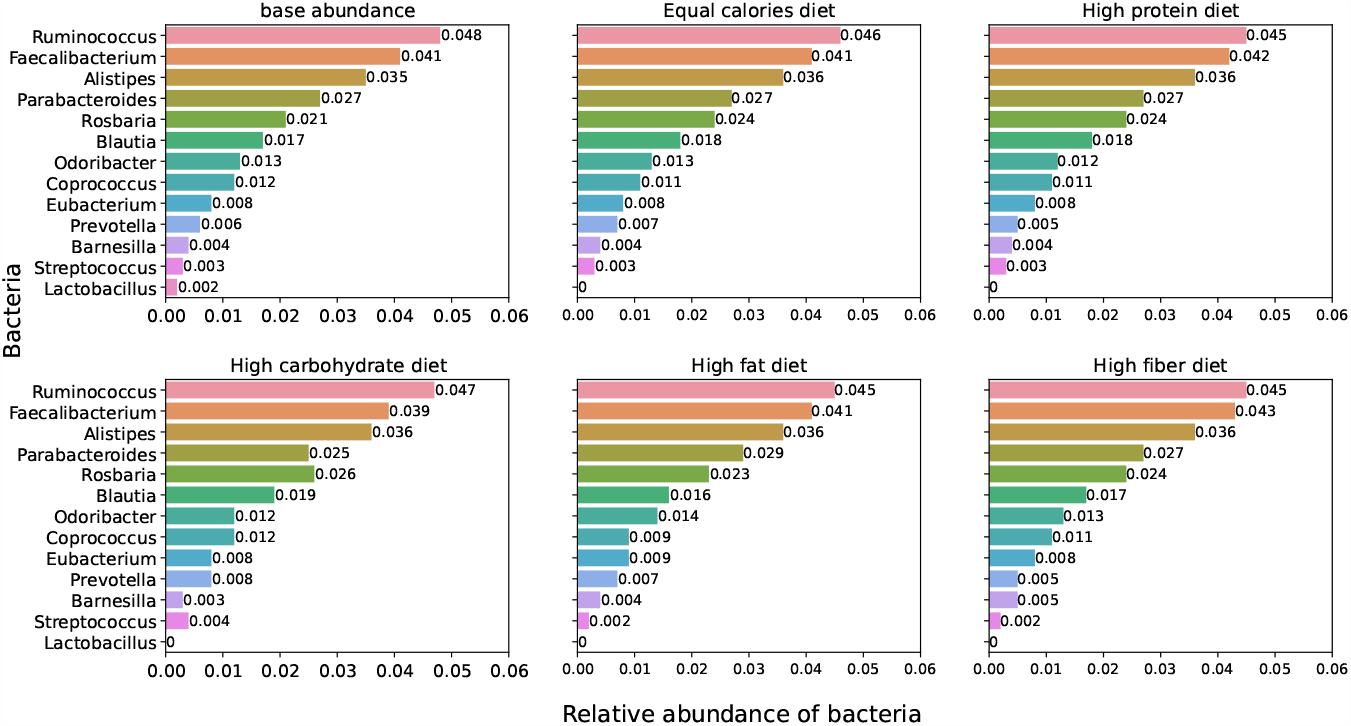
Relative abundance with different diets. A different representation of Figure 9 shows the change in the relative abundance of all bacterial strains except *Bacteroide* with a different diet.

#### 2.5.3 Relative population along the intestine

At present, there are only a few studies that monitor the change in the gut bacterial populations along the large intestine. These limited studies suggest *Bacteriorides, Ruminococcus* or *Firmicutes* and *Prevotella* in the proportion of 55%, 33%, and 12% in the ascending colon and 59%, 29% and 12% in the rectum [38]. In the 97 samples we examined and the 14 major bacterial types we simulated, there were no common occurrences of *Prevotella*. We compare the occurrence of the *Bacteriorides* and *Firmicutes*. As seen in Figure 8(b), these proportions vary along the intestine. However, the data is sparse for a direct comparison. More data will be helpful in learning or updating the model.

### 2.6 Scope of the work

Developing a comprehensive computational model which estimates the bacterial interactions with a growth model, with cross-sectional data available from cohorts rather than designing longitudinal studies, is ambitious. Clearly, the goal is ridden with several limitations. The number of bacterial strains included is large as a theoretical model, but nevertheless much less than the thousands of species. Extending the approach to such large-scale systems, if the data is available, will certainly require modifications to the algorithm. Similarly, the choice of four types of macronutrients may require adaptation. The statistical challenges of dealing with high dimensional parameter estimation with much lesser data is yet another challenge. However, with the various analyses we used, we show that the parameter estimation is relatively insensitive to the initial guesses. Given the complexity of the clinical data, and bacterial strains, the model is certainly at a model developmental stage. However, regardless of these limitations, the comprehensive approach presented in this work, from estimating the interaction parameters to comparing them with clinical observations, opens opportunities for various extensions. Other clinically relevant scenarios, such as the effect of periodic fasting use of drugs - antibiotic or non-antibiotic, can evolve as an extension of the current research.

## 3 Conclusion

In this work, we developed a comprehensive approach to model the 14 major types of gut bacteria. Towards the ultimate goal of having a model with clinical relevance, modeling different perturbations from the diet or drugs, we used readily available cross-sectional data of bacterial abundances to estimate the bacterial interaction parameters. Combining these estimates with the association with diet, several comparisons with the clinical data, such as the time to re-equilibrate after a perturbation or the association with diet, could be made.

## 4 Acknowledgements

We thank Prof. Kavita Jain for helpful discussions on modeling, and Dr. Benjamin Misselwitz for helpful discussions on gut microbiota. H.J. thanks CSIR for the JRF fellowship.

## 5 Additional Information

Data and codes are available at https://github.com/Himanshu535/-parameter_estimation

## 6 Supplementary Information

